# NMR Methods for Quantitative Isotopomer Rates in Real-Time Metabolism of Cells

**DOI:** 10.1101/544759

**Authors:** Michelle AC Reed, Jennie Roberts, Peter Gierth, Ēriks Kupče, Ulrich L Günther

## Abstract

Tracer-based metabolism is becoming increasingly important to study metabolic mechanisms in cells. NMR offers several approaches to measure label incorporation in metabolites, including ^13^C and ^1^H-detected spectra. The latter are generally more sensitive but quantification depends on the proton carbon ^1^*J*_CH_ coupling constant which varies significantly between different metabolites. It is therefore not possible to have one experiment optimised for all metabolites and quantification of ^1^H-edited spectra such as HSQCs requires precise knowledge of coupling constants. Increasing interest in tracer-based and metabolic flux analysis requires robust analyses with reasonably small acquisition times. Here we compare ^13^C-filtered and ^13^C-edited methods for quantification with a special focus towards application in real-time NMR of cancer cells under near-physiological conditions. We find an approach using a double-filter most suitable and sufficiently robust to reliably obtain ^13^C-incorporations from difference spectra. This is demonstrated for JJN3 multiple myeloma cells processing glucose over 24h.

## Introduction

Cellular metabolism changes extensively during normal proliferation and differentiation, and in response to disease and drug treatment. As the biological understanding of metabolism is vastly growing, there is also increasing interest in tracer-based experiments to study metabolic mechanisms and to carry out metabolic flux analysis. The two most common analytical technologies in this context are mass spectrometry (MS)^[1]^ and Nuclear Magnetic Resonance (NMR) spectroscopy. NMR is becoming an increasingly important method in this context^[2]^ as it has the ability to quantify site-specific label incorporation and is suitable for detecting metabolism in primary patient cells placed in an NMR tube under near-physiological conditions^[2,3]^.

NMR offers several options to measure ^13^C or ^15^N isotope incorporation. For low-concentration metabolism samples, ^13^C-observed spectra require long acquisition times because of the inherently lower sensitivity of ^13^C vs ^1^H. Moreover, long ^13^C relaxation times contribute to increased acquisition times, a factor that can be reduced using relaxation agents, although these substances are not desirable for use in cell cultures. For this reason, most studies of cell samples have used proton observed spectra. Fan and co-workers have successfully used TOCSY spectra to study label incorporation in metabolites^[4,5]^, but these are time-consuming and not suitable for high-throughput or real-time analyses, although recently developed fast acquisition schemes may soon overcome this limitation^[5]^. The most common approach for NMR based flux analysis is the use of ^1^H-^13^C-HSQC spectra, as originally suggested by Szyperski et al^[6–8]^. Several recent publications, including our own, show the true potential of HSQC spectra for a comprehensive tracer-based metabolic flux analysis^[9–11]^. However, intensities in HSQC spectra are unhelpfully modulated by the large variation of coupling constants in metabolites, which can range between 120 and 210 Hz. Fast acquisition schemes such as ALSOFAST considerably reduce acquisition times, however quantification is compromised due to the use of an Ernst angle approach^[12]^. There is considerable information content in CC coupling constants but this again requires long acquisition times^[9,11]^.

The use of simple proton observed 1D spectra has recently been proposed for high-throughput screening of label incorporation into metabolites^[13]^. In such 1D-^1^H spectra, the unlabelled ^12^C-species appears as a central peak, flanked by the ^13^C-coupled signals of the same proton. Vinaixa *et al*. suggested simply using the reduced intensity of the central signal to indicate label incorporation, but this approach is prone to large errors, most probably arising from signal overlap^[13]^. Alternatively, Wan et al^[14]^ suggested to calculate accurate concentrations from 1D-^1^H-^13^C-HSQC spectra using observed intensities and the pulse sequence specific transfer function. This is only possible if the ^1^*J*_CH_ coupling constant and relaxation rates are known which is unfortunately not the case for most metabolites. There have been earlier efforts to obtain quantitative HSQCs that cover a wide range of coupling constants. Q-HSQC uses a constant-time approach, which has the inherent disadvantage of lower sensitivity^[15]^.

Using a one-dimensional version of ^13^C-edited spectra can be very useful for samples with only a sparse set of metabolites. For such 1D-spectra, it is challenging to achieve artefact-free ^13^C-decoupling, and this is increasingly difficult at higher field strengths. In order to derive ^13^C/^12^C ratios it is also essential to acquire reference spectra that show all protons using a pulse sequence with equal transverse magnetization times in order to minimize relaxation-induced errors^[11]^. This allows intensities in spectra to be used directly without further scaling to calculate the proportions of ^1^H{^12^C} or ^1^H{^13^C}.

For one-dimensional approaches, there is a choice of using ^13^C-edited spectra, measuring all ^13^C-bound protons (^1^H{^13^C}) at a particular locus of a metabolite, or ^13^C-filtered spectra yielding the counterpart of protons attached to ^12^C only (^1^H{^12^C}). Such filtering is typically achieved using a BIRD sequence^[16]^. Our goal here is to quantify label incorporation in metabolic flux experiments using NMR methods in a manner that is suitable for high-throughput analyses or real-time NMR of living cells^[3]^. For this we have explored edited and filtered one-dimensional NMR methods. We have specifically looked at GBIRD^[17]^ as an option to quantify label incorporation in a range of metabolites with varying and unknown coupling constants.

## Results

We decided to compare what we saw as the most promising editing and filtering sequences and to look for the most robust option that works for a wide range of *J*_CH_ couplings, providing high-resolution data. Theoretically the difference between an all-proton experiment (^1^H{^12^C} + ^1^H{^13^C}) and a ^13^C-filtered experiment should yield the ^13^C-edited spectrum and vice versa, the ^13^C-edited subtracted from the all-proton experiment should yield the ^12^C-counterpart. This also requires that all spectra except the filtered are ^13^C-decoupled, although in practice it is advisable to also decouple the filtered spectrum in order to have exactly the same heat load arising from the decoupling sequence during acquisition thus avoiding difference artefacts.

For the editing experiments, we chose the recently published pulse sequence of Smith et al.^[11]^ (Figure S1A) as this sequence is already optimized for 1D-application and has a higher sensitivity than a 1D-HSQC. The chosen pulse sequence also allows a switch from editing ^1^H{^13^C} to all-^1^H by omitting INEPT-type ^13^C-pulses together with changes in the phase cycle. For filtering experiments, we implemented a double gradient-BIRD approach using two BIRD operators with delays 2*δ*=1/J_F1_ and 2*ε*=1/J_F2_ set for two different coupling constants^[16]^ (Figure S1B)^[17]^ to which we have added a bilevel adiabatic decoupling scheme. (See Supplementary Material and Figure S1 for technical details of this implementation). With this sequence, the corresponding all-^1^H experiment could be obtained by simply omitting selective ^13^C-pulses.

As an initial test sample we used a mixture of 48% [U-^13^C]glucose and 52% natural abundance glucose. The amount of label incorporation was confirmed using an undecoupled 1D spectrum by integrating the central ^1^H{^12^C} and outer ^1^H{^13^C} signals of the anomeric proton of glucose (Figure 1a). With decoupled spectra (Figure 1b-d and S2), the overall line shape with a resolution of 32k points for the all-proton spectra is complex as the signals of ^1^H{^12^C} and ^1^H{^13^C} are separated by a small 1.5-2Hz isotope shift. Filtered spectra yield narrower lines because the ^1^H{^12^C} resonance is not subject to ^13^C-mediated relaxation.

**Figure 1.**
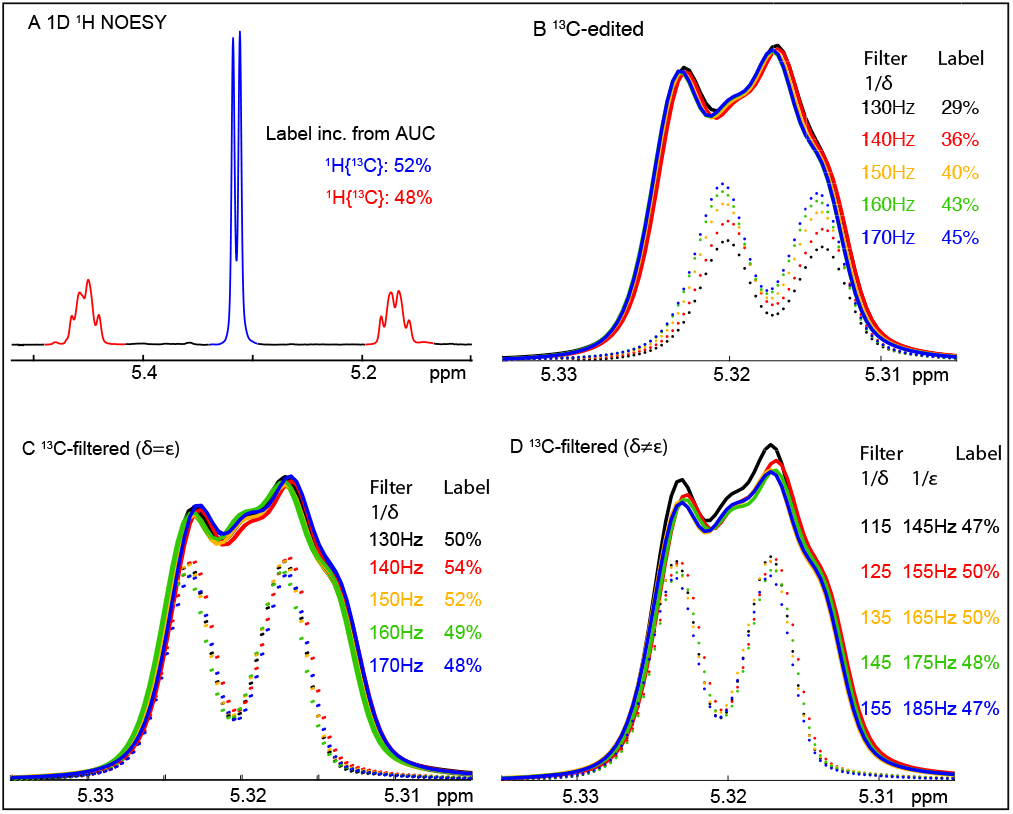
Editing and filtering efficiencies. Extracts from 1D spectra showing the anomeric ^1^H from α-D-glucose (^1^*J*_CH_=168Hz) for a 48% U-^13^C-glucose and 52% natural abundance glucose in RPMI cell culture medium. (A) 1D ^1^H-NOESY spectrum with the ^1^H{^12^C} and ^1^H{^13^C} resonances highlighted in blue and red respectively. (B) Overlay of 1D ‘all-^1^H’ (solid lines) and 1D ^1^H{^13^C}-edited (dotted lines) spectra showing reduced accuracy as the *J*_F_ value determining the evolution periods (2*δ*=2*ε*=1/*J*_F_) deviates increasingly from the optimal value of 168Hz. (C) and (D): Overlays of 1D ‘all-^1^H’ (solid lines) and 1D ^1^H{^12^C} ^13^C-filtered (dotted lines) spectra showing the efficiency of the double-BIRD filter with two identical filter periods (C) or with two different filter periods (D).

To address the question of how effective the filtering approach is for sub-optimal evolution periods, we arrayed the filtering periods without ^13^C-decoupling during acquisition to observe residual ^1^H{^13^C} resonances. This eliminates potential errors in intensities arising from suboptimal ^13^C-decoupling. For this we focussed on the well-resolved resonances of the anomeric ^1^H of α-D-glucose with a *J*_CH_ coupling constant of 170Hz using the double filter GBIRD with either both filter periods *δ* and *ε* set to the same value (Figure 2, blue line and Figure S3a), or with the two filter periods set to *J* values separated by 50Hz (Figure 2, black line and Figure S3b).

**Figure 2.**
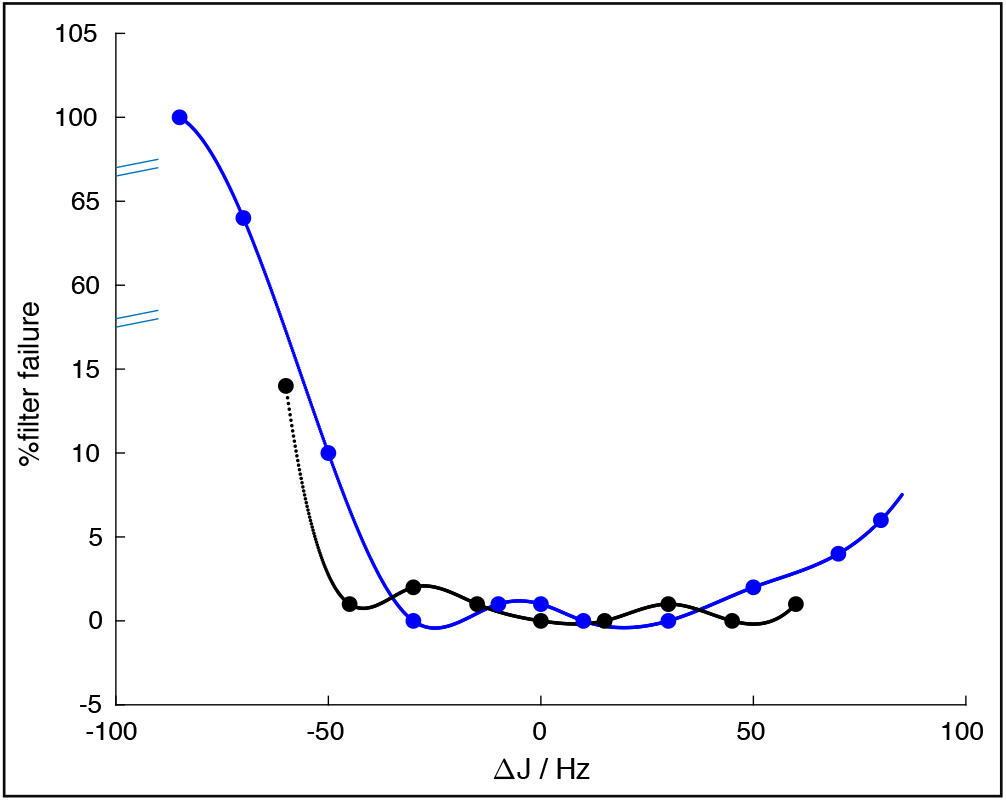
Filter failure, the proportion of ^1^H{^13^C} resonance intensity of the anomeric ^1^H from α-D-glucose that survives the filter in undecoupled ^13^C-filtered spectra. Double GBIRD filter with two equal filter periods 2*δ*=2*ε*=1/*J*_filt_ (blue line) and double GBIRD filter with two unequal filter periods, 2*δ*=1/*J*_F1_ and 2*ε*=1/*J*_F2_, separated by 50Hz (black line). Filter failures were calculated from the areas under the curve for the ^1^H{^13^C} resonances as 100*(AUC/AUC_J85Hz_).

For the experiments with both delays set to the same value, the filter efficiency suffers for large offsets of these delays from the actual coupling constant. For two unequal delays *δ* and *ε* set 50Hz apart, the filter becomes considerably more tolerant towards suboptimal settings, in fact, the ^1^H{^13^C} resonance always appears to be well suppressed. The filter is somewhat more robust towards delays set for larger *J* values than smaller. Generally, the double filter GBIRD with two different filter periods separated by 50Hz is a very robust filter, well suited to isotopomer quantification in metabolites.

With decoupling during acquisition, the situation is somewhat more complicated, mainly owing to the isotope shift that leads to two sets of overlapping doublets. Figure 1a shows the overall ^1^H-NOESY signal without decoupling, whereas panels b-d show experiments with decoupling. The results again illustrate the greater robustness of the double filtering approach. As expected, editing shows a variation of signal intensity for varying settings of the evolution delay owing to the transfer function. The filtering experiments proved much more robust, although with decoupling there is a less obvious difference in the quality of filtering for equal and different BIRD periods with non-ideal evolution constants (Figure 1c and Figure 1d).

A remaining overestimation of the proportion of ^1^H{^12^C} in decoupled experiments is observed for longer filtering delays (smaller *J*_F_). This was puzzling as the undecoupled experiment (Figure 2 and S3) indicated that the filter efficiency was very high. The likely explanation relates to the faster relaxation of ^1^H{^13^C} compared with ^1^H{^12^C}. For smaller *J*_F_, i.e. longer *δ* and *ε*, the transverse magnetization period is increased. The all-^1^H resonance intensity is more affected by transverse relaxation than the ^1^H{^12^C} and so for lower values of *J*_F_, there is a small overestimate ofthe proportion of ^1^H{^12^C} present. The faster relaxation of ^1^H{^13^C} vs ^1^H{^12^C} is clearly reflected in the greater line widths (Figure 1B-D). Nonetheless, the filtering approach is much more robust to non-ideal *J* than the editing approach.

We tested the filtering approach in real-time NMR spectra using JJN3 multiple myeloma cells fed with [U-^13^C]glucose as a metabolic precursor. For the acquisition of these spectra, JJN3 cells were embedded into a matrix of agarose as previously described^[3]^. As real-time spectra change over time, the pulse sequence had to be modified to acquire the filtered and all-proton reference spectrum in a scan interleaved mode. Decoupling was applied for both spectra in order to have the same heating effect for both, even though the ^13^C-filtered spectrum does not benefit from decoupling. The acquisition time for each pair of spectra was 15m for a sample containing approximately half a million cells. The time course as shown in Figure 3 shows the difference spectra of all-^1^H and ^1^H{^12^C}spectra, demonstrating the suitability of this approach for real time acquisition of metabolic changes using a tracer-based approach.

**Figure 3.**
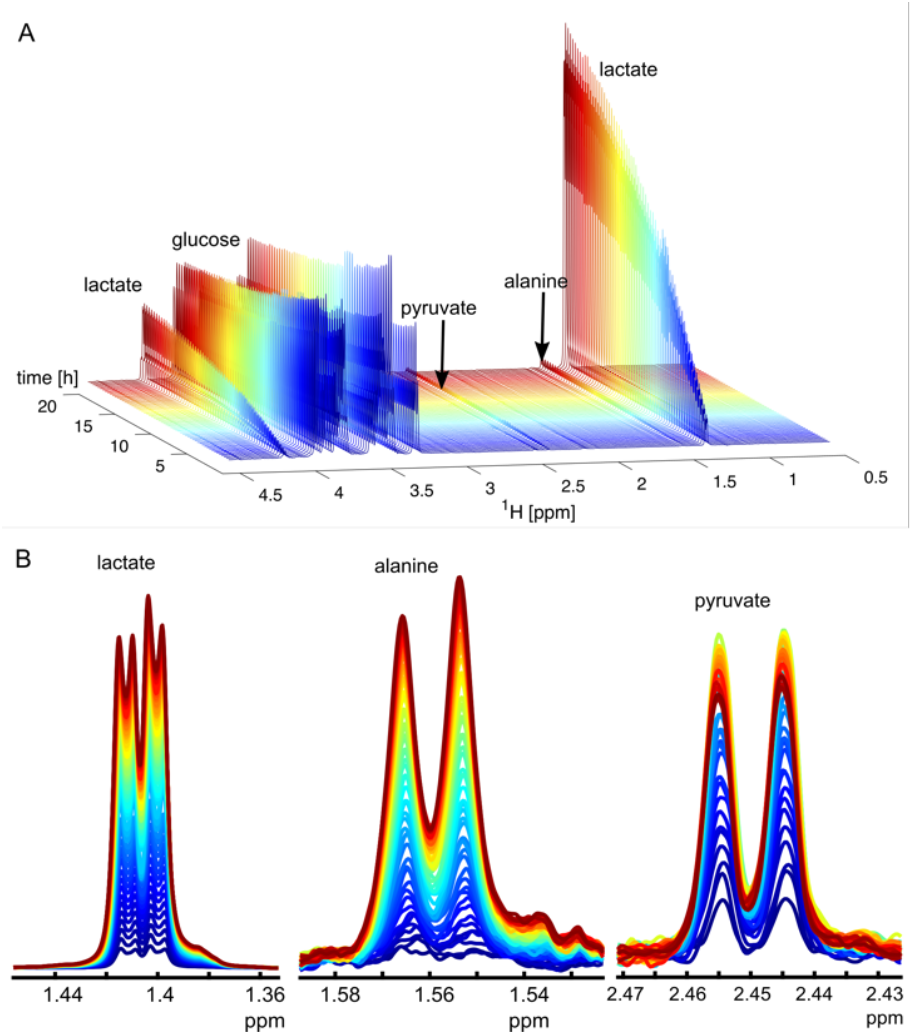
Real-time experiment with JJN3 cells. Spectra were acquired using the double GBIRD filter with *δ*=1/128Hz and *ε*=1/168Hz. ‘All ^1^H’ and ^1^H{^12^C} spectra were acquired in scan-interleaved mode. The spectra shown, representing ^1^H{^13^C}, were calculated by taking the difference between all-^1^H and ^1^H{^12^C} spectra. As the decoupling bandwidth did not cover the region around 200ppm we did not decouple lactate carboxylic acid carbons, hence the lactate CH_3_ signal (B) shows the extra ^3^J_CH_ coupling from COO to the methyl protons (further experimental details in Supplementary Material).

It should be noted that decoupling restricts the number of points that can be acquired. At 600MHz and a spectral width of 12ppm, 32K points could be acquired on both room temperature and 5mm cryoprobes. The room temperature probe is less able to compensate for heating effects of adiabatic decoupling and, therefore compromises were made as detailed in the methods section. We have also used this approach using a 1.7mm cryoprobe at 600MHz which has a superconducting ^1^H receiver coil that is more sensitive to heating. Nevertheless, good quality spectra could be obtained for 16k data points.

The ^13^C-filtered approach is particularly useful to measure label incorporation in metabolites with unknown and varying *J*_CH_ coupling constants. The maximum filter leakage is 2% when the filter periods are mis-set by +/−50Hz from the ideal values for a particular ^1^H{^13^C} resonance.

When coupling constants are known, editing becomes a more realistic option for the quantification of label incorporations as the overall transfer function can be calculated based on the ^1^*J*_CH_ and *T*_2_ transverse relaxation times, as recently shown by Wan et al. for 1D-HMQC^[14]^. However, this requires precise knowledge of these parameters. Moreover, for real-time approaches, where the same metabolites are compared over a longer time period, the filter approach yields highly reproducible spectra with a good baseline and water suppression, without losses incurred by echo-anti-echo selection.

In summary, we have shown that label incorporation in sparsely labelled metabolism samples can be obtained using ^13^C-filtered one-dimensional spectra using a sequence with two filter constants that is highly tolerant towards deviations of the set delays from the actual coupling constant. With this approach, label incorporations can be obtained with a relatively high accuracy. This method requires the availability of isolated signals in a 1D spectrum, although such filters could be combined with twodimensional spectra to reduce signal overlap.

## Materials and Methods

### Sample preparation

The glucose sample was in RPMI cell culture medium with 10% D_2_O. The 25mM glucose was a 50:50 mixture of nominally 98% [U-^13^C]glucose and natural abundance glucose.

For real-time NMR, JJN3 cells were maintained at exponential growth phase in RPMI1640 bicarbonate-buffered cell culture medium supplemented with 10% foetal calf serum in a humidified incubator at 37°C and 5% CO_2_. For NMR experiments, approximately half a million cells were resuspended in a 0.1% w/v agarose/RPMI matrix with 10% D2O and 1mM sodium 3-(trimethylsilyl)propionate-2,2,3,3-d4. 600μl of cell suspension was loaded into a standard 5mm NMR tube and transported to HWB.NMR at 37°C.

### NMR Experiments

All experiments were performed at 310K on Bruker AVANCE II or III spectrometers equipped with 5mm probes. For all simple ^1^H 1D experiments, high resolution was achieved by acquiring 32K data points for an acquisition time of ca. 2s with adiabatic ^13^C decoupling using Wurst decoupling pulses in the bilev p5m4 supercycle (bi_p5m4sp_4sp.2). A recycle time of 7.5s was used with a total of 40 dummy and actual scans, giving an experiment time of *ca*. 5min. The ^13^C carrier frequency was set to 80ppm and the chirp π pulses were used throughout.

Initial experiments used a 11.7T magnet with a room temperature TXI probe. A room temperature probe is not ideal for experiments with long ^13^C-decoupling periods as there is substantial sample heating. To reduce this, the air flow rate was increased from 400 to 550lph. To reduce the power going into the probe due to ^13^C-decoupling during acquisition, the adiabaticity factor of the pulses was reduced from Q=5 to 2.5 with pulses generated using the wavemaker with the shape, cawurst-20 (160ppm, 0.7 or 1.4ms; Q=2.5). 24 dummy scans were required for the sample temperature to stabilise sufficiently for good resonance linewidths. The spectral width was 15.971ppm/7978.7Hz and all proton pulses were at 12.9W.

For the ^13^C-filtering experiments, gradient pulses 1 to 4 were 1ms pulses with gradient strengths of 21.4, −21.1, 35.8 and 21.9 G/cm respectively. For the editing experiments, gradient pulses 1 and 2 were 2.5ms pulses at 45.5 and 46.5 G/cm respectively. The INEPT gradient pulses were 1ms 34.2 G/cm pulses. Further experiments were performed at 310K on a 14.1T magnet with a cold TXI probe. The experiments were acquired as above except that it was not necessary to increase the air flow rate and only 8 dummy scans were required for the sample temperature to stabilise. Therefore, 32 scans were acquired with just 8 dummy scans. The spectral width was 11.986ppm/7183.9Hz and all proton pulses were at 5.5W.

For the filtering experiments, gradient pulses 1 to 4 were 1ms pulses with gradient strengths of 15.0, - 21.1, 35.8 and 21.9 G/cm respectively. For the editing experiments, gradient pulses 1 and 2 were 2.5ms pulses at 10.2 and 5.4 G/cm respectively. The INEPT gradient pulses were 1ms 16.1 G/cm pulses.

### Real-time experiments

A time-course of *ca*. 19 hours was acquired using a 14.1 Tesla magnet equipped with a cold TXI probe and an AVANCE II console. Initially, the time-course was recorded by repeating a one-hour loop of 4 experiments 16 times. Each loop consisted of the filtered and edited experiments with their corresponding reference experiments. Parameters were as described above for cold probes except that the number of scans was increased to 96 giving an individual experiment time of 15mins.

The attempt to subtract the ^1^H{-^12^C} spectrum from the corresponding all-^1^H spectrum to obtain a difference spectrum with resonances from ^1^H{-^13^C} gave a spectrum with many artefacts due to changing shim and particularly changing pH due to metabolism by the JJN3 cells in the NMR tube. Therefore, a pulse program was created to run the ^1^H{-^12^C} and all-^1^H experiments in scan-interleaved mode. This gave two free induction decays and the difference spectrum gave the residual signal attributable to ^1^H{^13^C} with greatly reduced artefacts. Memory limitations on this console necessitated a reduction in TD to 16K because of the two memory buffers required for the difference experiment (This is a specific problem on some older BRUKER AVANCE III consoles under Topspin3.6). To partly offset the resulting decrease in acquisition and to reduce the loss in resolution in the resulting spectra, the spectral width was reduced to 5995.2Hz/10ppm, giving a 1.3s acquisition time. The number of dummy and actual scans was set to 16 and 60 respectively giving an experiment time of approximately 15min.

Decoupling for the real-time spectra shown in Figure 3 was achieved using Wurst decoupling pulses in the bilev p5m4 supercycle with a decoupling bandwidth of 160ppm, centred around 80ppm (at 600MHz proton frequency). At this bandwidth we do not decouple lactate carboxylic acid carbons, hence the lactate CH_3_ signal (B) shows the extra ^3^*J*_CH_ coupling from COO to the methyl protons. Spectra were acquired with a spectral width of 10ppm and 16k points giving an acquisition time of 1.36sec. The relaxation delay was 5.3s and spectra were acquired with 60 scans (and 16 dummy scans) for each subspectrum, giving a 15min difference experiment. These difference spectra give a clean account of the label incorporation into individual metabolites.

## Acknowledgements

We thank BBSRC for supporting JR in the context of a BBSRC CASE studentship. We are also grateful to the Wellcome Trust for supporting access to NMR instruments at the Henry Wellcome Building for Biomolecular NMR in Birmingham.

## Supplementary Figures

**Figure S1:**
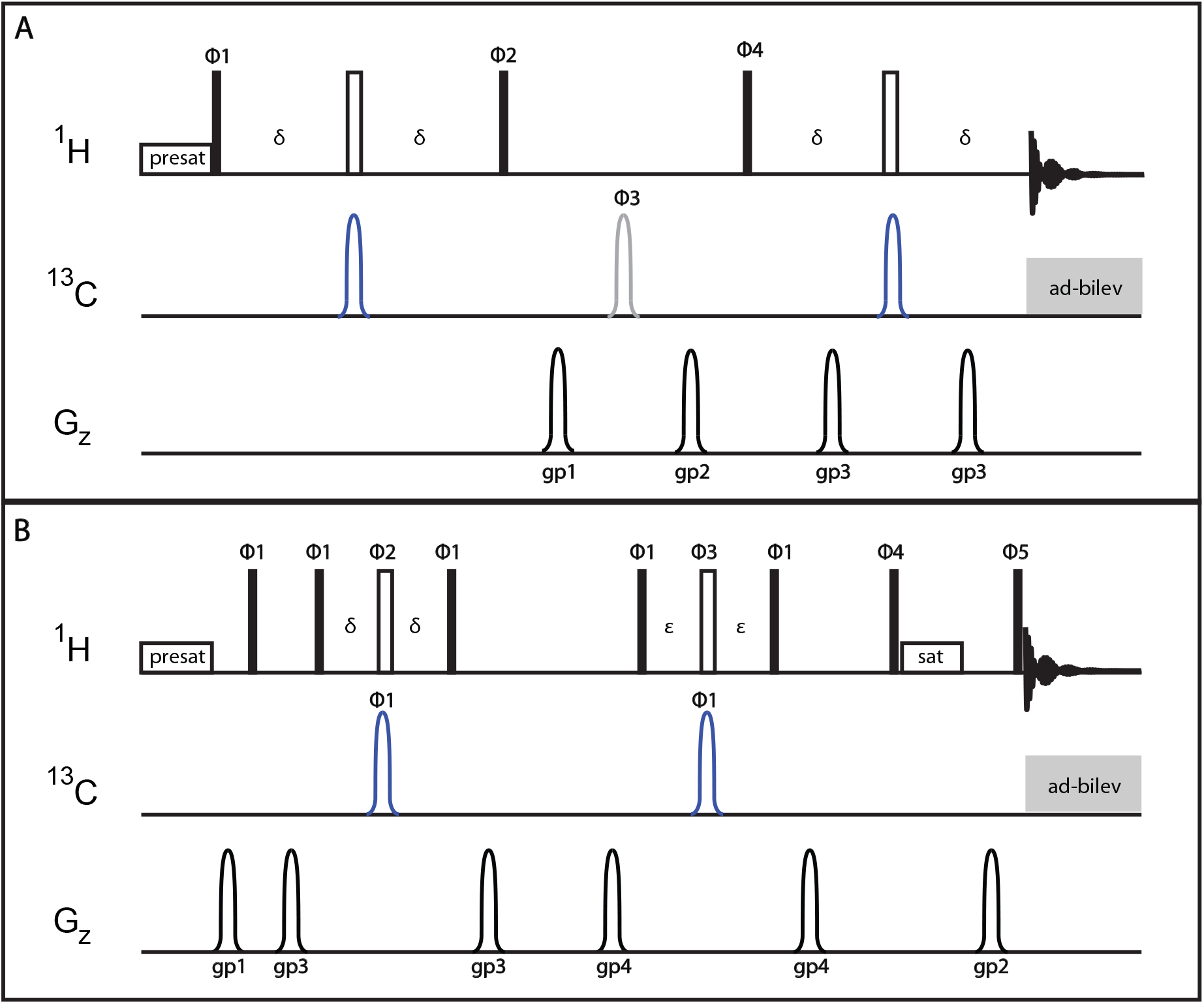
^13^C-editing and filtering pulse sequences. **A**: ^13^C-editing sequence of Smith et al. included here for clarity. After the first INEPT block (δ=1/4J), ^1^H{^13^C} magnetisation is longitudinal whereas ^1^H{^12^C} magnetisation is transverse and is destroyed by the gradients, gp1 and gp2. The grey ^13^C pulse, which is only executed every second transient, improves pathway selection in combination with changes in the phase cycles of ϕ3, ϕ4 and the receiver. Phase cycles were as follows: ϕ1=0 0 2 2, ϕ2=1 1 3 3, ϕ3=0 1 2 3, ϕ4=1 3 3 1 2 0 0 2 and receiver phase=0 2 2 0 3 1 1 3. The pulse sequence used for the reference non-selective experiment was identical except that the blue ^13^C pulses were omitted and the phase cycle changed slightly with ϕ2=0 0 2 2, ϕ3=0 and ϕ4=0 2 2 0 1 3 3 1. **B**: Double BIRD-^13^C-filtered 1D ^1^H NOESY. The initial ^1^H pulse puts ^1^H{^12^C} and ^1^H{^13^C} magnetisation transverse. ^1^H{^12^C} magnetisation changes sign as it passes through each of the BIRD elements. Consequently, it is defocussed and then refocussed by the matching gradients either side of each BIRD element. ^1^H{^13^C} magnetisation does not change sign and so is destroyed by the gradients. The delays δ and ε were set to 1/2J_1_ and 1/2J_2_ respectively where J_1_ and J_2_ are the two ^1^J_CH_ couplings targeted for filtering. Phase cycles were as follows: ϕ1=0 2, ϕ2=1 1 3 3, ϕ3= 3 3 1 1, ϕ4=0 0 0 0 0 0 0 0 2 2 2 2 2 2 2 2, ϕ5=0 0 2 2 1 1 3 3, and receiver phase=0 2 2 0 1 3 3 1 2 0 0 2 3 1 1 3. The pulse sequence used for the reference non-selective experiment was identical except that the power of the ^13^C pulses was zero. Thus ^1^H{^12^C} and ^1^H{^13^C} magnetisation passes through the sequence in the same way. In all sequences, ^13^C π-pulses were chirp pulses. To obtain high resolution, 32K points were acquired, necessitating a long acquisition time (~2s). Decoupling during this period used Wurst decoupling pulses in the bilev p5m4 supercycle. Water suppression is fairly efficient in the selective experiments because of the gradients. However, slightly better water suppression is achieved with the GBIRD filter sequence.

**Figure S2:**
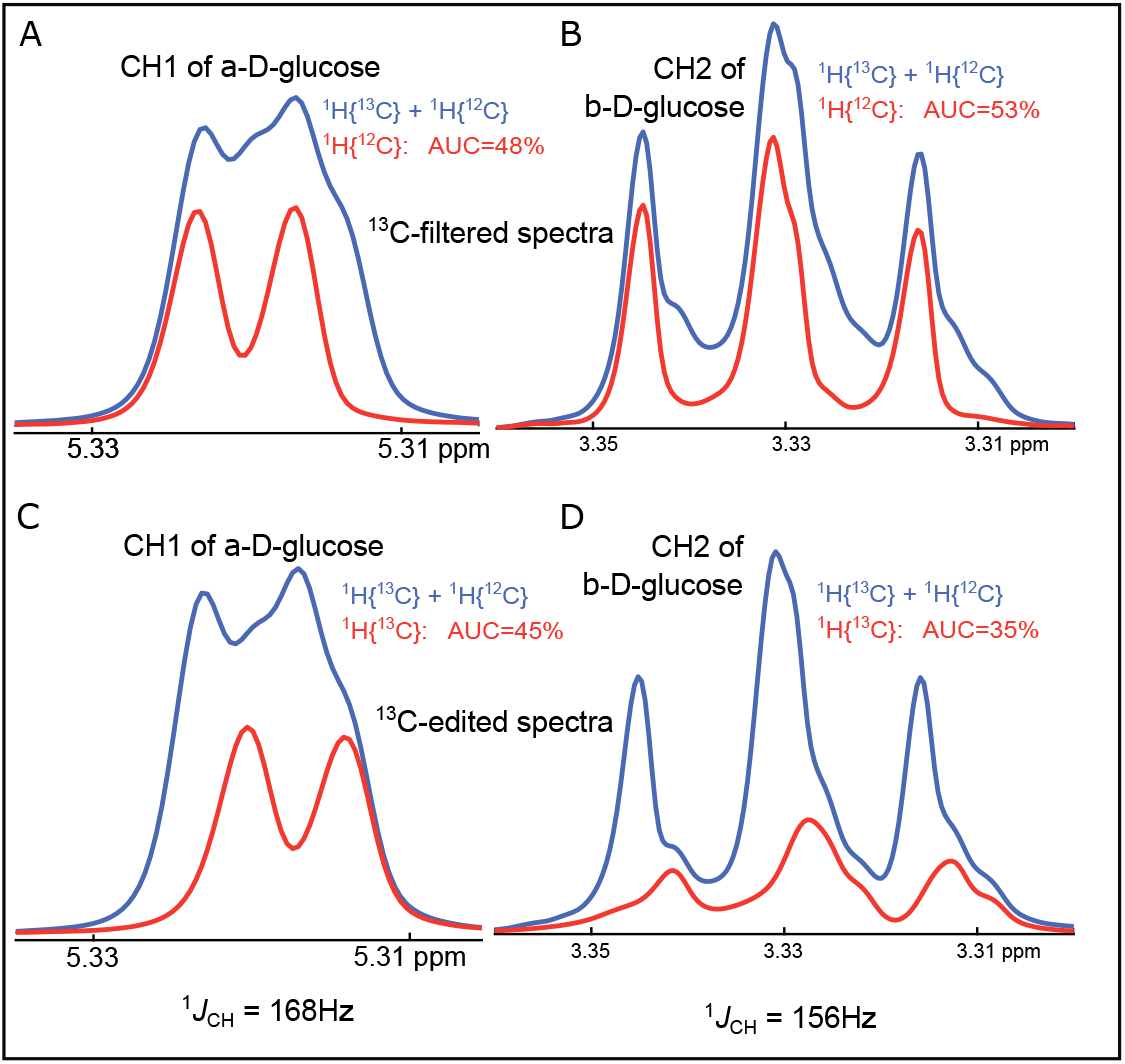
Regions from 1D spectra of 48% U-^13^C-glucose and 52% natural abundance glucose in RPMI cell culture medium. A&B: Overlays of 1D ‘all-^1^H’ no-BIRD (blue) and 1D ^1^H{^12^C} double BIRD-filtered NOESY (red) for the anomeric ^1^H1 of α-D-glucose region (A) and the ^1^H2 of β-D-glucose (B) respectively. C&D: Overlays of 1D ‘all-^1^H’ (blue) and 1D ^1^H{^13^C}-edited (red) spectra for the anomeric ^1^H1 of α-D-glucose region (C) and the ^1^H2 of β-D-glucose (D) respectively. Complex line shapes arise from a small isotope upfield shift of 1.5-2Hz observed for ^1^H{^13^C} in the edited spectrum compared with ^1^H{^12^C} in the filtered spectrum and because filtered spectra (red lines in Fig.S2a and S2b) yield narrower lines because the ^1^H{^12^C} resonance is not subject to ^13^C-mediated relaxation. The scalar coupling evolution periods were set for *J*=170Hz for the ^13^C-edited experiment, and 170Hz and 140Hz for double-filtered BIRD. Spectra were acquired with adiabatic ^13^C decoupling to minimize sample heating. For the editing experiments, the evolution periods *δ* were set to 1/*J* with *J*=170Hz whereas for the GBIRD double-filtering experiments, the two filter delays were set to *δ*=1/*J*_F1_ and *ε*=1/*J*_F2_ with *J*_F1_=170Hz and *J*_F2_=140Hz. Because of the different linewidths of ^12^C and ^13^C associated protons and the isotope shift, it was necessary to integrate the area under the curve (AUC) to estimate the proportion of ^1^H{^12^C} and ^1^H{^13^C}. AUCs indicate that filtering and editing are both very effective for the anomeric ^1^H of α-D-glucose (*J*=168Hz) with estimated proportions of ^1^H{^13^C} and ^1^H{^12^C} of 45% (actual 48%) and 48% (actual 52%), respectively. However, for the H2 of β-D-glucose with a different ^1^*J*_CH_ coupling constant of 156Hz, the estimated proportions are 35% (actual 48%) for the editing, and 53% (actual 52%) for the filtering experiment, showing a large error for the editing experiment arising from a suboptimal *δ* delay. Here the advantage of the double filter becomes evident.

**Figure S3:**
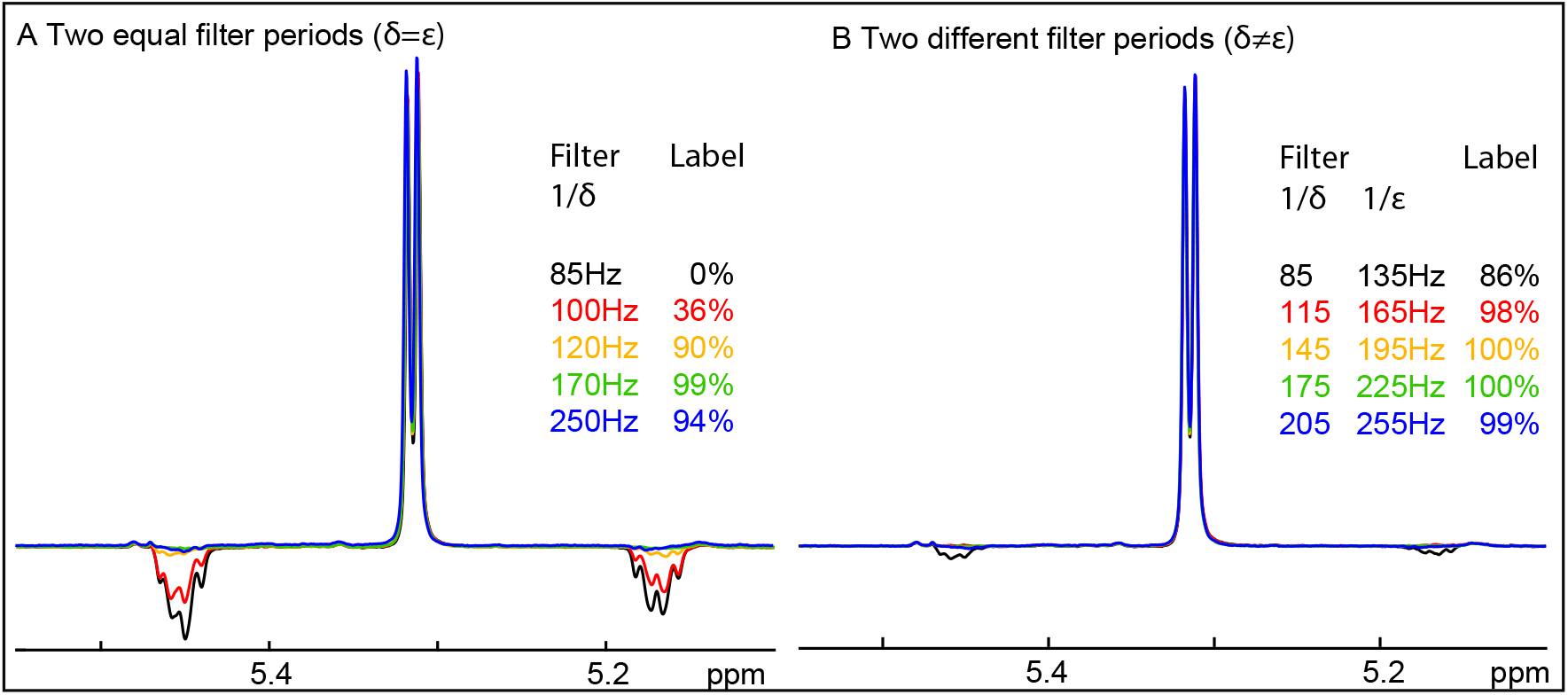
Filter efficiency for the resonances of the anomeric ^1^H from α-D-glucose. Extracts from 1D spectra of 48% U-^13^C-glucose and 52% natural abundance glucose in DMEM media. a) Double GBIRD filter with two equal filter periods *δ*=*ε*=1/J_filt_. b) Double GBIRD filter with two unequal filter periods, *δ*= 1/J_F1_ and *ε*=1/J_F2_. Filter efficiencies were calculated from the areas under the curve for the ^1^H{^13^C} resonances, as 100*(1-AUC/AUC_J85Hz_).

